# Genomic Insights of Bruneian Malays

**DOI:** 10.1101/2022.06.01.492266

**Authors:** Mirza Azmi, Lie Chen, Adi Idris, Zen H. Lu

**Affiliations:** PAPRSB Institute of Health Sciences, Universiti Brunei Darussalam, Jalan Tungku Link, BE1410 Brunei Darussalam; Griffith Centre for Cell and Gene Therapy, Menzies Health Institute Queensland and School of Pharmacy and Medical Science, Griffith University, Southport, QLD 4222, Australia

## Abstract

The Malays and their many sub-ethnic groups collectively make up one of the largest population groups in Southeast Asia. However, their genomes, especially those from Brunei, remain very much underrepresented and understudied. Here, we analysed the publicly available WGS and genotyping data of two and 39 Bruneian Malay individuals, respectively. NGS reads from the two individuals were first mapped against the GRCh38 human reference genome and their variants called. Of the total ∼5.28 million short nucleotide variants and indels identified, ∼217K of them were found to be novel; with some predicted to be deleterious and associated with risk factors of common non-communicable diseases in Brunei. Unmapped reads were next mapped against the recently reported novel Chinese and Japanese genomic contigs and *de novo* assembled. ∼227 Kbp genomic sequences missing in GRCh38 and a partial open reading frame encoding a potential novel small zinc finger protein were successfully discovered. Interestingly, although the Malays in Brunei, Singapore and Malaysia share >83% common variants, principal component and admixture analysis comparing the genetic structure of the local Malays against other Asian population groups suggested that they are genetically closer to some Filipino ethnic groups than the Malays in Malaysia and Singapore. Taken together, our work provides the first comprehensive insight into the genomes of the Bruneian Malay population.

## Introduction

The Malays are an Austronesian ethnic group consisting of various sub-ethnic populations residing mainly in the Southeast Asian countries of Brunei, Malaysia, Singapore and Indonesia. Despite them being one of the largest ethnic groups in the region, Malay genomes were not included in either the earlier HapMap or the later 1000 Genomes Project (1KGP) [1, 2]. In fact, to date, only a handful of genomic studies have been conducted in Singapore, Malaysia and Indonesia [3–10]. Although valuable information in areas such as population genetic structure, rare or novel genetic variations, pharmacogenomics and genetic disease risk factors have been gained, much remain to be uncovered. This is especially true for the different Malay sub-ethnic groups in Brunei where their genomics landscape remained virtually unexplored till now.

With a population of about 430 thousand, Brunei is located on the northern coastline of Borneo and it is the smallest country by population in Southeast Asia. Approximately 65% of the population are Malays, which politically, historically, and culturally are made up of seven indigenous groups namely Melayu Brunei, Melayu Tutong, Melayu Belait, Kedayan, Murut, Dusun, and Bisaya. The last three are also found throughout the island of Borneo; including the neighbouring countries of Malaysia and Indonesia. Studies have suggested that they are likely to be genetically related to the Amis in Taiwan who might have migrated to Borneo through the Philippines [6, 10, 11].

Furthermore, recent studies have reported genomic sequences which are unique to their specific East Asian populations but missing in the human reference genome [12–16]. Not only have these sequences provided better insights into the population genetic structure between the Southern and Northern Han Chinese, for example, they have also allowed identification of several novel spliced variant gene transcripts. Therefore, the existence of novel sequences in the Malay genome is not totally unlikely and together with the genetic variants identified from the various Malay sub-ethnic groups, they should provide a more accurate genetic stratification of the Malay populations.

In this study, the recently published whole genome sequencing (WGS) and genotyping data of several Bruneian Malays were reanalysed [6, 10]. Short variants as well as copy number variations (CNVs) were called and compared with those reported in other Malay groups in the region. Population genetic structure analysis was also conducted by comparing a subset of the single nucleotide variants (SNVs) against several Asian population groups to provide better understanding on the genetic architecture of the Bruneian Malays. Finally, attempts were made to search for novel genomic sequences within the Bruneian Malay genomes. Findings from our study have not only shown, for the first time, differences in genetic ancestry between Malays in Brunei and Southeast Asian regions but also discovered novel sequences of the Bruneian Malay.

## Materials and Methods

### Subjects

Genomic data of 41 Bruneian Malays used in this study were obtained from two published studies; namely WGS data of two Dusun female individuals generated as part of the Simons Genome Diversity Project, and 730K genotyping data of 20 Dusun and 17 Murut individuals, respectively, from a Southeast Asian population structure study [6, 10] (Table S1). In addition, the 700K AncestryDNA^®^’s genotyping data of a local mixed-race Malay-European mother-son duo (private contribution) was also included in the population structure analysis.

### Variant calling and annotation

The WGS and genotyping data were analysed using a two-pronged approach (Figure S1). The quality-assessed NGS reads were first mapped against the GRCh38 human reference genome with decoy sequences [17] using the maximal exact matches algorithm of BWA (version 0.7.17) [18]. In addition to the default mapping options, insert size was set to 400 bp and shorter split and unpaired paired-end reads were both tagged as secondary mapping.

Three different variant callers; namely GATK (version 4.1.4.1) [19], BCFtools (version 1.9) [20], and FreeBayes (version 1.0.2) [21] were then employed to call both SNVs and insertions/deletions (INDELs) from the mapped genomes. The respective GATK’s “best practices” workflows for identifying somatic and mitochondrial variants were adopted. In brief, the mapped reads were name- or position-sorted, mates fixed, duplicates marked, and base calls recalibrated against known variant datasets obtained from 1000 Genomes Project [2], dbSNP (build 154), Singapore Genome Variation Project (SGVP) [3] and Singapore Sequencing Malay Project (SSMP) [4]. Coordinates of the Singaporean variants were up-lifted from GRCh37 to GRCh38 using GATK’s “LiftoverVcf” command. Nuclear variants from each of the two recalibrated BAM files were next called using the GATK’s “HaplotypeCaller” and jointly genotyped using “GenotypeGVCFs”. Quality scores of these variants were subjected to further recalibration against the same known variant datasets and also those obtained from GRCh38-lifted-over HapMap. Variants which did not fit the quality statistics were discarded.

When calling short variants using BCFtools, the “mpileup” command was first used to calculate the likelihood of genotypes at specific positions with the minimum mapping quality and number of INDEL-supporting gapped reads set to 30 and 5, respectively. Raw variants were next generated using the “call” command with the multi-allelic calling algorithm switched on and the genotype quality (GQ) and posterior probabilities (GP) calculated before filtering away low-quality ones (QUAL<Q30, GQ<Q30, and DP<10×).

Short variants on reads with mapping quality ≥Q30 were also called from the two BAM files using another haplotype-based variant caller, FreeBayes, running on default parameters. BCFtools was then used to filter the raw variants as previously described.

To minimise the limitations of each of the three variant callers [22, 23], a consensus set of variants was generated by intersecting the three filtered VCF files using the “isec” command of the BCFtools. In addition, INDELs longer than 40 bp were also removed to further improve the confidence of the consensus variants.

On the other hand, variants of the other 37 Bruneian Malays were extracted from a PLINK-formatted genotyping dataset and transformed into a VCF-formatted file using the program PLINK (version 1.9) [24]. A custom script was used to transform the variant tables from AncestryDNA^®^ to their respective VCF-formatted files. GRCh37-based genomic coordinates of all variants were finally lifted as described previously.

All variants called from the 41 Malays were next merged using BCFtools before being annotated against genomic features in various databases (Table S2) using the “annotate” command of BCFtools and AnnoVar [25]. Features added included associated dbSNP’s accession number; ENSEMBL’s gene transcripts; GnomAD’s allelic frequency of variants; GWAS’s and ClinVar’s clinical association (*e.g.* genetic diseases or pharmacogenomics); and potential functional impacts of non-synonymous variants from SIFT and PolyPhen-2. Biological and molecular functions of genes with potential deleterious variants, which were defined as non-synonymous coding variants annotated as ‘deleterious’ by both SIFT and PolyPhen-2 or classified as ‘high impact’ by ENSEMBL, were investigated by submitting them to the Gene Ontology online web-server (http://geneontology.org/). On the other hand, mitochondrial non-synonymous variants were uploaded to the MITOMAP web-server (https://www.mitomap.org/) [26] and their functional impacts predicted using the APOGEE [27] function available there.

In addition, basic variant statistics on allele frequency, depth distribution, Het/Homo and Ts/Tv ratios were calculated using the statistical function of either BCFtools or RTG tool [28]. Variant density distributed across each chromosome was calculated based on a 1 Mbp sliding window using the SNPdensity command of VCFtool [29] and plotted using the CMplot R package [30].

### CNV Calling

cn.MOPs was employed to identify CNVs in the two WGS datasets [31]. Read depths within a 1-Kbp sliding window were first calculated using the “getReadCountsfromBAM” function across each of the 22 autosomes before they were normalised and plotted with the “segplot” function. CNVs meeting the following four criteria, namely (1) read depth ≥10×; (2) spanning across three or more segments/windows; (3) log2 value for copy number gain ≥0.8; and (4) log2 value for copy number loss ≤-2.8; were then extracted and annotated using the AnnotSV (https://www.lbgi.fr/AnnotSV/) web-server [32].

### Uncovering Novel Sequences in the Bruneian Malay Genome

NGS reads from the two Bruneian Malays which failed to be mapped to the human reference genome were subjected to further investigation to search for novel sequences that may be unique to the Bruneian Malay genome. It is probable that some of these unmapped reads were in fact of microbial origins since salivary DNAs were used in the WGS. Therefore, two rounds of microbial mapping were applied to remove these microbial reads (Figure S2A). Firstly, they were BWA-mapped, as previously described, against the Human Oral Microbiome Database (HOMD) (http://www.homd.org/ftp/all_oral_genomes/current/). Next, converted unmapped fastq reads from the first round mapping were fed into Kraken2 metagenomics analysis package [33] to further fish out as many remaining microbial reads as possible by comparing them against the MiniKraken2 (ftp://ftp.ccb.jhu.edu/pub/data/kraken2dbs/old/minikraken2_v1_8GB_201904.tgz) microbial database.

The presence of potential novel sequences in Bruneian Malay genomes was finally investigated by subjecting the final set of unmapped reads to a workflow involving mapping against East Asian sequences and *de-novo* assembly (Figure S2B). A novel sequence is defined here as one that is missing from the human reference genome.

The recently reported ∼12 and ∼6 Mbp novel HX1 Chinese [12] and JRG Japanese [15] sequences, respectively, were chosen as the “reference genomes” for the mapping as comparative genomic analysis here showed that they are genetically the closest population groups, among those publicly available, to the Bruneian Malays. Again, a similar BWA mapping strategy used earlier was applied. However, the output BAM files were more stringently filtered (>Q30 mapping quality, >30× coverage depth, and mapped region spanning over the length of a read (*i.e.* >100 bp)) to increase the confidence that the mapped reads were indeed similar to the Chinese/Japanese sequences. Variant calling and filtering was next carried out as before using BCFtools’ “mpileup”/”call” and “filter” (QUAL and GQ ≥30) commands. Consensus sequences representative of the Bruneian Malay were then generated using GATK’s “FastaAlternateReferenceMaker” which replaced the reference alleles with the filtered variants.

Next, *de novo* assembly of the unmapped reads was performed using a k-mer-based assembler, MEGAHIT [34]. k-mer values ranging between 17 and 31 (as recommended by a best k-mer predictor, KmerGenie [35]) with an increment of 2 (parameter: --k-min 17 --k-max 31 --k-step 2) were attempted. MEGAHIT by default removes redundant contigs by merging those sharing ≥95% similarity and trims edges with low-coverage (<4×). The newly assembled contigs were then BLASTed with default parameters against the GRCh38 human reference genome to identify those of human origin. Sequences were considered human when they shared ≥90% identity with GRCh38. Those with ≤90% sequence identity were BLASTed one more time but against the NCBI’s non-redundant (NR) nucleotide database. This time, only contigs with ≥95% identity to human fosmids- and BAC-cloned human sequences were considered likely novel Bruneian Malay sequences.

Contigs derived from the mapping and *de novo* assembly were next BLASTed against each other to remove redundant sequences. The final set of human contigs were further investigated for their gene coding potential by BLASTing their 6-frame translations against the online NCBI’s non-redundant protein database.

### Comparative Genomic Analysis

The Malay variant data files of (1) 89 and 25 genotyped individuals from SGVP and Morseburg’s study, respectively, and (2) 96 whole genome sequenced individuals from SSMP were transformed, when necessary, and merged as previously described into two separate VCF files according to the country of origin. These were then compared against variants of the 41 Bruneian Malays using the “isec” command of BCFtools to identify shared and unique variants among the Malays in the three countries. Potential functional impacts of those that were unique to the Bruneian Malays were further investigated as previously described.

To gain more insights into the population genetic structure of the Malays in Brunei, a principal component (PCA) and an admixture analysis were performed on a subset of SNP from 1,499 individuals, including the 41 Bruneians, belonging to 22 different South, East, and Southeast Asian population groups (Table S3). As a control, the AncestryDNA^®^’s genotyping variants of a European relative of the two mixed-race Bruneians were included. Again, whenever necessary, SNP files were transformed into GRCh38-based VCF files as previously described.

The PCA was conducted according to a method adopted from a previous study investigating the European population genetic structure [36]. Prior to calculating the principal components of these SNPs, a filtering step using PLINK was performed to increase the examining resolution of the SNPs while avoiding the removal of excessive number of true positives, *i.e*., SNPs with low linkage disequilibrium (LD). The filtering criteria included (1) missingness rate of ≥10%, *i.e*., below 90% genotyping rate to ensure all the SNPs were comparable across all the different population groups; and (2) genotypic *r^2^*, which is a common measurement for LD, ≥80% within a 50-SNP sliding window increasing at 5 SNPs per step to ensure the selected SNPs were all independent from one another as well as non-arbitrary. The first ten principal components of each individual with the filtered SNPs were then measured using the PLINK’s command “pca” and those which could explain over 80% of the variance were selected and plotted against each other using the R package, ggplot2 [37].

An “unsupervised” admixture analysis on the same filtered SNP datasets was next conducted using the default settings of ADMIXTURE [38]. The analysis assumed no prior knowledge on ancestral origin and historical context of the observed genotypic profiles. The number of major ancestral origins (K) in each of the Asian population groups were first estimated and cross-validated before the hypothetical ancestral proportions (Q) of each of the 1,499 individuals were calculated at K=4, 5 and 6. A bar plot representing their hypothetical ancestral proportions was subsequently plotted using ggplot2.

## Results

### Genomic landscape of the Bruneian Malays

The high-quality WGS paired-end reads from each of the two Bruneians could be mapped successfully across >94% the human reference genome at an average depth of 37x and 47x, respectively (Table S4). The remaining ∼6% falling mainly within known gaps or challenging chromosomal regions, such as centromere and telomeres (Figure S3).

Of the ∼5.07 million consensus variants called from the two Bruneian Malays, ∼217K, consisting mainly of INDELs, were novel and have not been reported in dbSNP (Table 1). Both Het/Homo and Ts/Tv ratios were within the range reported in other Asian populations, indicating the reliability of the variant calling [4, 5, 9]. Interestingly, the two individuals were found to harbour ∼150K rare variants with GnomAD’s minor allele frequency (MAF) >0.001.

**Table 1.** Statistics of the consensus variants called from the WGS data of the two Bruneian Malays

| Variants | Combined | Individual 1 | Individual 2 |
| --- | --- | --- | --- |
| **<u>Nuclear Chromosomes</u>** |  |  |  |
| Total: | 5,069,437 | 3,935,796 | 3,935,869 |
| SNPs (% total) | 4,569,044 (90.4%) | 3,551,224 (90.0%) | 3,544,852 (89.9%) |
| INDELs (% total) | 500,393(9.6%) | 384,572 (10.0%) | 391,017 (10.1%) |
| Het/Homo ratio | 1.23 | 1.25 | 1.22 |
| SNP Ts/Tv ratio | 2.09 | 2.09 | 2.09 |
| Novel variants (% total): | 216,837 (4.3%) | 161,427 (4.1%) | 167,745 (4.3%) |
| SNPs (% novel) | 7,565 (3.5%) | 4,468 (2.8%) | 5,497 (3.8%) |
| INDELs (% novel) | 209,272 (96.5) | 156,959 (97.2%) | 162,248 (96.2%) |
| MAF <0.001 (% total) | 148,481 (2.9%) | 77,660 (2.0%) | 80,143 (2.0%) |
| RNA genes* (% total) | 614,368 (12.1%) | 477,073 (12.1%) | 477,184 (12.1%) |
| Protein-coding genes* (% total): | 1,910,236 (37.7%) | 1,478,330 (37.6) | 1,478,164 (37.6) |
| Intron (% coding genes) | 1,743,454 (34.4%) | 1,349,176 (34.3%) | 1,348,714 (34.3%) |
| Exon (% coding genes) | 29,591 (0.6%) | 22,865 (0.6%) | 22,770 (0.6%) |
| 5' UTR (% coding genes) | 9,700 (0.2%) | 7,526 (0.2%) | 7,624 (0.2%) |
| 3' UTR (% coding genes) | 42,748 (0.8%) | 33,332 (0.8%) | 33,095 (0.8%) |
| Splicing (% coding genes) | 190 (<0.01%) | 143 (<0.01%) | 147 (<0.01%) |

\*Variants in the primary assembly of GRCh38 only

Both SNPs and INDELs were found to be distributed fairly evenly throughout the whole genome with an overall mean density of 1,498 SNPs/Mbp and 145 INDELs/Mbp, respectively, for the two local individuals (Figure 1). A total of 22 and 20 SNP- and INDEL-dense regions, *i.e.* ≥2x mean density, respectively, could be identified (Table S5 & S6) and perhaps, not unexpected, the human leucocyte antigen (*HLA*) locus (chr6:29-33Mbp) was among one such regions. However, there were only two other SNP-dense regions (chr6:5-9Mb & chr16:77-79Mb), which have been reported elsewhere [4, 39].

**Figure 1A.**
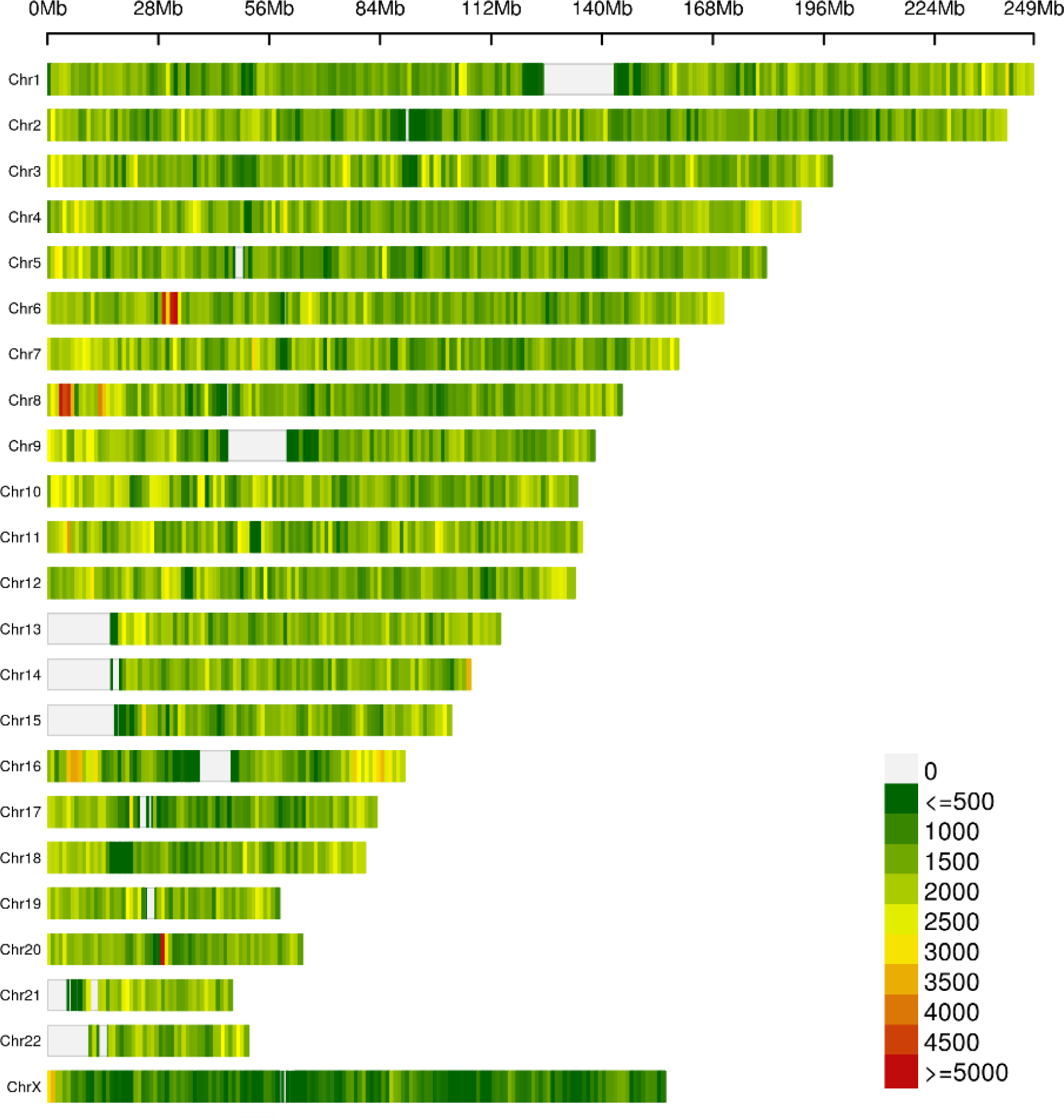
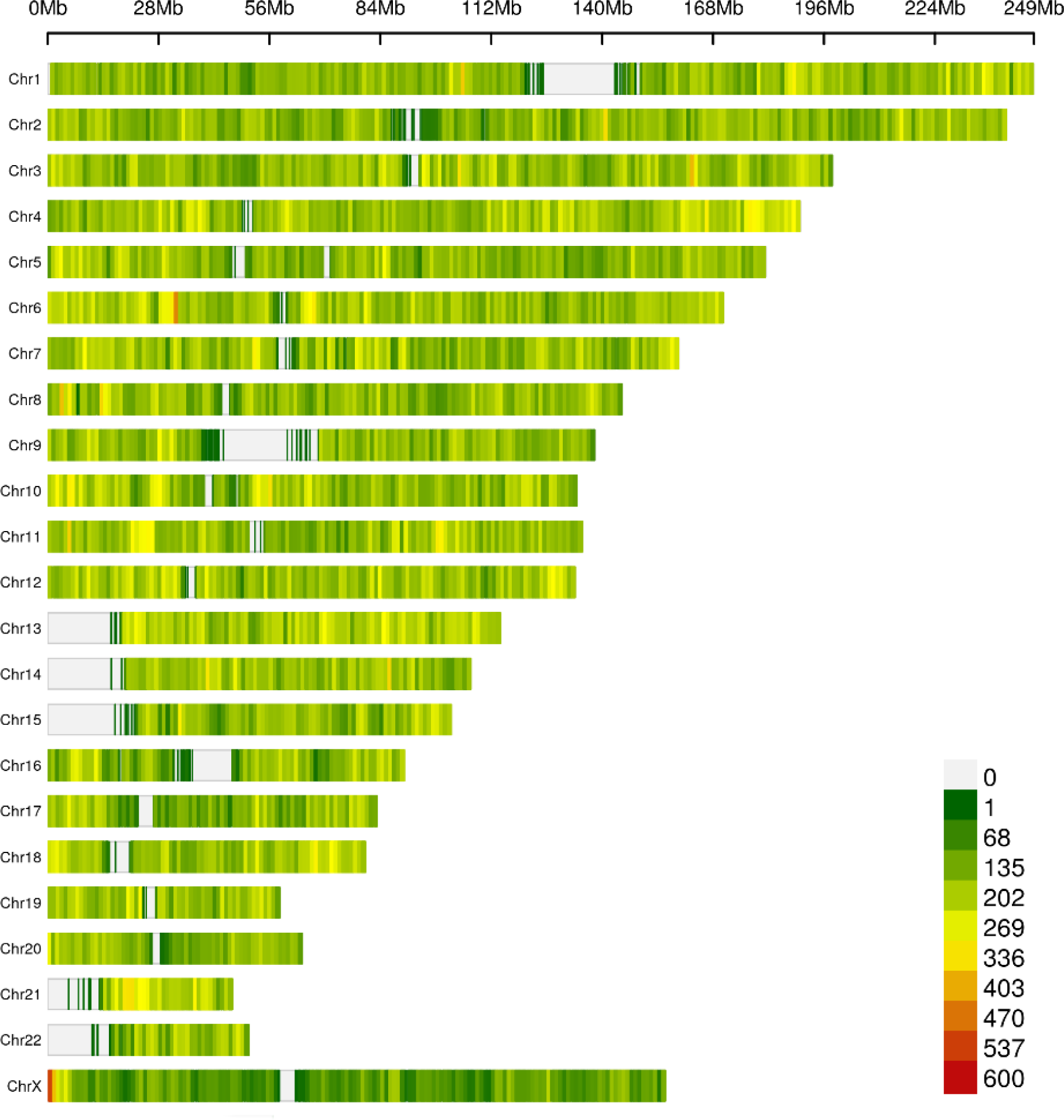
Variant density plot of the two Malay genomes. The SNPs (A) and INDELs (B) were distributed across the entire genome except in the unmapped regions and known gaps of the human reference genome. Regions with high variant densities are coloured red while those in green depict regions with low variant densities.

Furthermore, as a source of genetic diversity among different population groups, the number of CNVs found on the 22 autosomes of the two Bruneian Malays were estimated to total 1,192. The 392 and 859 copy number gains and losses, respectively; ranged in sizes from the minimum 3 Kbp set by the caller to the longest 159 Kbp. 62 of them were found to span over 72 different protein-coding genes and pseudogenes and they could potentially disrupt the associated gene structure. In fact, a number of these genes are known to be associated with diseases such as cancer and cardiovascular diseases (Table S7).

### Novel Sequences in the Bruneian Malay Genome

A total of ∼93 million unmapped or ∼3% of the raw reads from the two WGS datasets were obtained after initial mapping against the human reference and microbial genomes. Of these, ∼16 million were mapped to ∼146 Kbp sequences making up of ∼9.2 and ∼137 Kbp novel Chinese and Japanese contigs, respectively. 170 short variants, including both SNPs and INDELs, were called from the mapping; suggesting the potential commonality of these sequences among the three population groups.

Furthermore, although more than 19K contigs with a cumulative length of ∼53 Mbp could be assembled from the “leftover” reads, only 58 of them totalling ∼82 Kbp were found to share >95% BLAST search sequence identity with either some regions of the GRCh38 human genome or fosmids- and BAC-cloned human sequences, *i.e.,* they are of human origin. The “discarded” contigs were likely to belong to some other yet-to-be verified human or unculturable microbial or misassembled sequences.

The mapping and *de-novo* assembly approaches had, therefore, yielded 227,763 bp novel Bruneian sequences which are missing in the GRCh38 human reference genome. When the gene-coding potential of these sequences were further investigated, an open reading frame out of a total 473 was found to share 92% sequence identity across a 106-residue segment with a macaque’s hypothetical small zinc finger protein (Figure S4). The protein is predicted to be involved in the homeostasis of zinc ions.

### Comparative genomics among the Malays

A total of 5,276,758 short variants consisting of ∼4,8 million SNPs and ∼0.5 million INDELs were obtained from the 41 Bruneian Malays when ∼700k genotyping variants were merged with the ∼5.07 million WGS variants (Table 2). Of these, ∼36K equal number of synonymous and non-synonymous variants were found to fall on coding regions of genes. Among the latter, 2,094 were predicted to be deleterious with the potential to impact 1,718 genes. Gene Ontology analysis revealed that these genes are involved in cellular processes such as immune defence, protein modification, transcriptional regulation, cell signalling and molecular transport (Figure 2). However, it is important to note that majority of these deleterious variant may have no clinical manifestation. In fact, those with minor allele frequencies <0.05 were found in only small number among the 41 Malay individuals (Table 3). Such variants are predicted to have a higher likelihood of being the risk alleles of diseases [40]. Indeed, some of these risk alleles are associated with common non-communicable diseases, *e.g.* breast cancer and heart disease, in Brunei. When compared with the publicly available variants of Singaporean and Malaysian Malays, 83% (∼4.38 million) of the Bruneian variants could be found within the ∼14.02 million Singaporean dataset while 94.6% (∼0.61 million) of the ∼0.65 million Malaysian variants intersected with that of the Bruneian; *i.e.* they are genetically highly similar to one another. However, the variants shared by the three Malay groups are expected to be higher if larger and more representative cohorts from Malaysia and Brunei are available.

**Figure 2.**
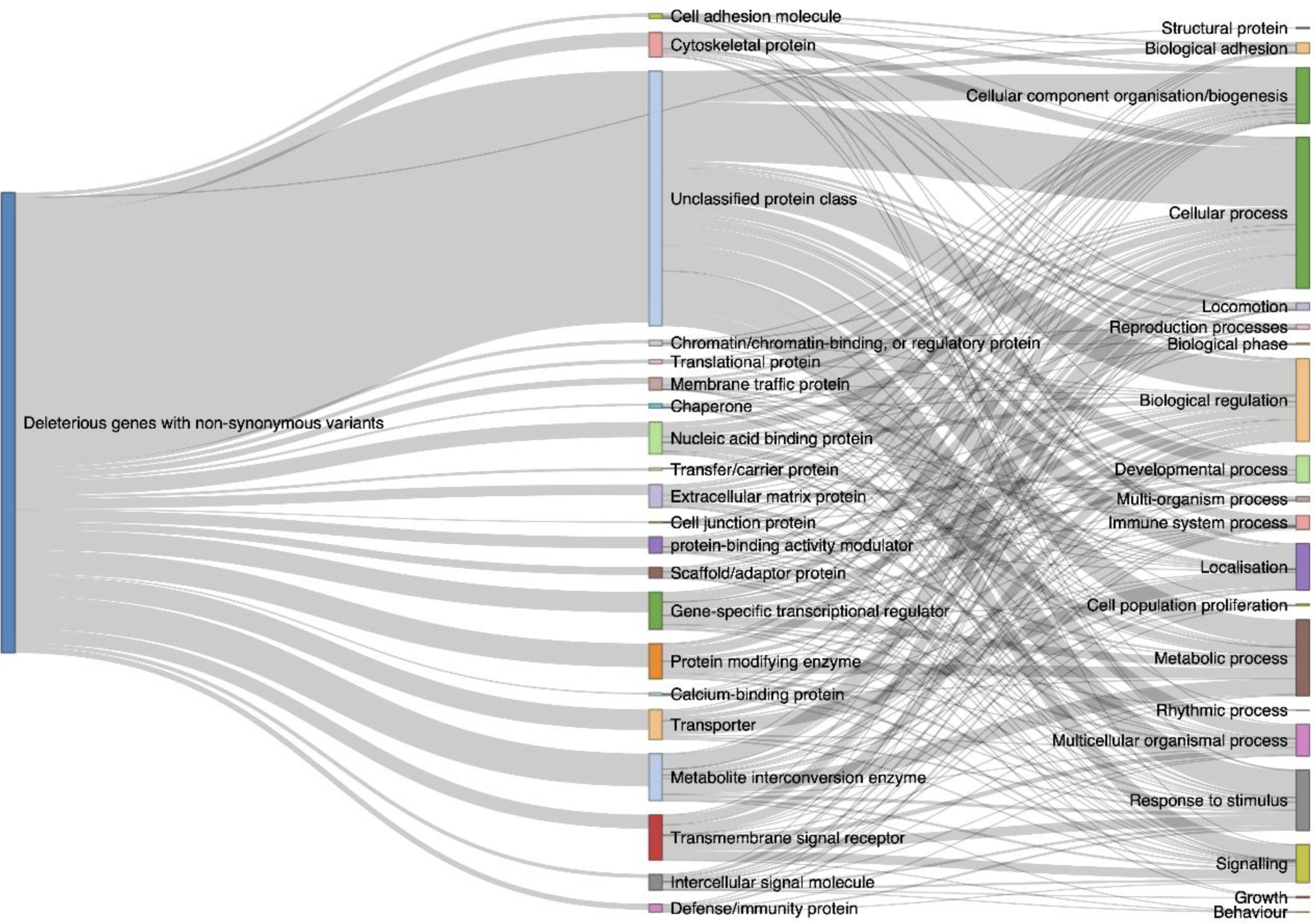
Gene Ontology analysis on genes harbouring deleterious variants. The Sankey map shows the association between genes and their protein classes on the middle and biological functions on the right.

**Table 2.** Summary of coding variants found in the 41 Bruneian Malays

| **Functional annotation** | **Combined** | **WGS** | **Genotyping** |
| --- | --- | --- | --- |
| Short variants: | 5,276,758 | 5,069,437 | 697,223 |
| *SNP* | 4,776,365 | 4,569,044 | 697,223 |
| *INDELs* | 500,393 | 500,393 | 0 |
| Protein-coding variants: | 35,783 | 29,591 | 17,952 |
| *Synonymous (%)* | 17,900 (50.0%) | 14,811 (50.1%) | 9,242 (51.5%) |
| *Non-synonymous (%)*: | 17,753 (49.6%) | 14,652 (49.5%) | 8,690 (48.4%) |
| *Missense (%)* | 17,111 (96.4%) | 14,039 (95.8%) | 8,611 (99.1%) |
| *Stop-gain SNPs (%)* | 172 (1.0%) | 146 (1.0%) | 66 (0.8%) |
| *Stop-loss SNPs (%)* | 35 (0.2%) | 32 (0.2%) | 13 (0.1%) |
| *Frameshift INDELs (%)* | 154 (0.8%) | 154 (1.1%) | 0 (<0.1%) |
| *Non-frameshift INDELs (%)* | 281 (1.6%) | 281 (1.9%) | 0 (<0.1%) |
| *Unclassified (%)* | 130 (0.4%) | 128 (0.4%) | 20 (0.1%) |

**Table 3.** Potential risk allele of diseases identified in the local Malays

| dbSNP ID | Gene | ENSEMBL Transcript | Mutation | Global AF<sup>§</sup> | EAS AF<sup>§</sup> | BN AF<sup>§</sup> | Associated risk factors | Reference |
| --- | --- | --- | --- | --- | --- | --- | --- | --- |
| rs5174 | *LRP8* | ENST00000306052 | c.G2855A:<br>p.R952Q | 0.28 | 0.03 | 0.027 | Myocardial infarction | [45] |
| rs2075291 | *APOA5* | ENST00000542499 | c.G553T:<br>p.G185C | 0.0027 | 0.0713 | 0.013 | Hypertriglyceridemia | [46] |
| rs766173 | *BRCA2* | ENST00000380152 | c.A865C:<br>p.N289H | 0.0385 | 0.0932 | 0.041 | Breast cancer | [47] |
| rs1801133 | *MTHFR* | ENST00000376585 | c.C788T:<br>p.A263V | 0.2719 | 0.2873 | 0.041 | Increased drug toxicity of cyclophosphamide and fluorouracil (cancer) | [48] |
| rs4148323 | *UGT1A1* | ENST00000305208 | c.G211A:<br>p.G71R | 0.0092 | 0.1419 | 0.03 | Increased drug toxicity to deferasirox (iron overload) | [49] |

<sup>§</sup> Allele Frequencies (AF) were based derived from GnomAD

### Distinct admixed genetic structure of the Bruneian Malays

To gain a better understanding on the population genetics of Bruneian Malays in the context of the wider Asian population groups, *the first ten* principal components (PC) of 583,453 SNPs from each of the 1,498 East, South, and Southeast Asian individuals and a control individual with European ancestry were calculated. A two-dimension PCA plot of the first two PCs, which together could explain over 80% of the variance, shows that PC1 separated the South Asians from the other population groups while PC2 separated the overlapping East and Southeast Asians (Figure 3A). The extreme outlier on PC1 *was* the control European individual. The genetically less varied and, hence, the packed South Asian cluster included various population groups residing in the Indian Subcontinent and other countries. On the other hand, the North-South overlapping between the tightly clustered East Asian groups and the more spread-out cluster of the Southeast Asian*s* indicated a probable flow of genetic materials among the different groups in the region.

**Figure 3.**
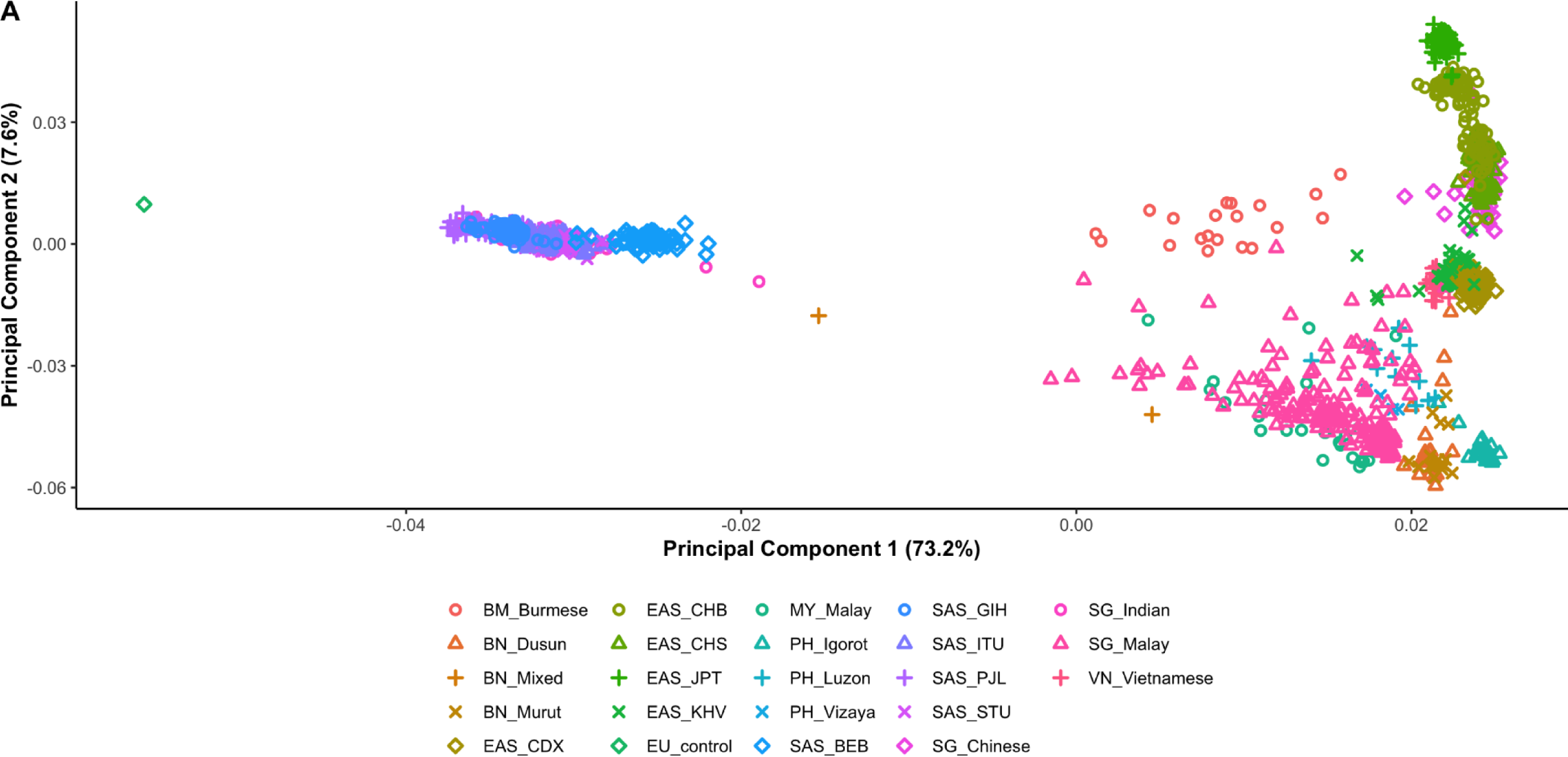
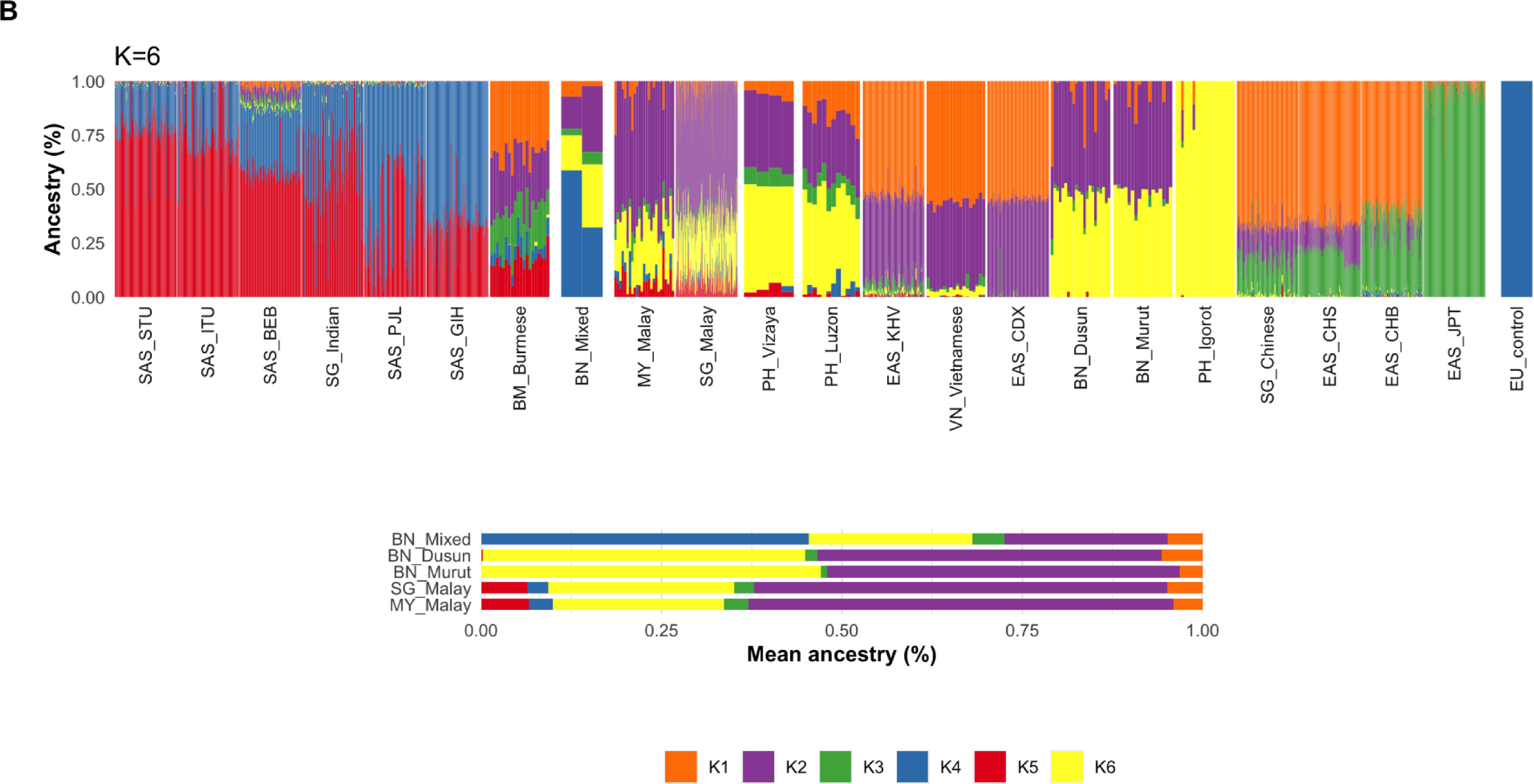
Population genetic structure of 22 Asian population groups. (A) Principal component analysis and (B) admixture analysis of ∼583K variants among the 1499 individuals. BM, Burma/Myanmar; BN, Brunei; EAS, East Asians; CDX, Chinese Dai; CHB, Han Chinese in Beijing; CHS, Southern Han Chinese; JPT, Japanese; KHV, Kinh Vietnamese, EU, Europe; MY, Malaysia; PH, Philippines; SAS, South Asians; BEB, Bengali; GIH, Gujarati; ITU, Telugu; PJL, Punjabi; STU, Sri Lankan Tamil; SG, Singapore; VN, Vietnam.

Although culturally and linguistically the most similar among all the Malay groups in Southeast Asia, subtle genetic differences among the Malays in Brunei, Singapore, and Malaysia could be further inferred from their PCA clustering. Not surprisingly, the two Bruneian Malay sub-ethnic groups, the Murut and the Dusun, are genetically highly similar to each other as they compactly crowded together within the Southeast Asian cluster. On the other hand, the Singaporean and Malaysian Malays, which themselves formed another compact subcluster, are slightly further apart from the Bruneian Malays. Regardless, the two Malays are not that dissimilar from each other. In addition, the Malays from Brunei, Singapore, and Malaysia are also genetically closely related to the two Filipino ethnic groups, namely Luzon and Vizaya. Many of them overlapped with one another in the PCA plot. Interestingly, as one of the oldest indigenous groups in the Philippines, the Igorots themselves formed another compact subcluster which lied furthest away from the other Southeast Asians.

An “unsupervised” admixture analysis was then conducted under the assumption that the 22 Asian populations are made up of four to six major ancestral groups, *i.e*., K=4, 5 and 6. Of the three K values, K=5 exhibited the lowest cross-validation error, though the difference between each K values was mostly negligible. Therefore, the hypothetical ancestral proportions (Q) of each of the 1,499 individuals, including the control, were first calculated with all three K values. At K=4, differences in the ancestral components between majority of the South Asians and the European control were almost indistinguishable and separation among the different population groups became clearer only at K=5 and above (data not shown). While K=5 has the lowest cross-validation error, K=6 appears to be in best agreement with findings from the PCA (Figure 3B).

While K2 constituted the major ancestral components among all the Malay population groups in Brunei, Singapore, and Malaysia, the subtle genetic differences observed in the PCA segregation became clearer in the admixture analysis. The genetic ancestry of the Bruneian Malays is made up largely of two components of K2 and K6, amounting to 91% and 95% in the Dusun and Murut, respectively, and small proportions of K1 and K3. In contrast, majority of the Singaporean and Malaysian Malays share a highly similar genetic admixture pattern containing all six ancestral components, including the European K4 and South Asian K5 ancestries. Unlike their Bruneian cousins, the K2 component of the Singaporean and Malaysian Malays was found to be present in a higher (>57%) proportion while K6 contributed only 26% of the admixed make-up. In fact, the result here suggested that Bruneian Malays share a closer genetic ancestry background with the Vizaya and Luzon from Philippines. It is, therefore, clear that although the three Malay population groups in Southeast Asia may share an almost identical culture and language, differences do exist in their genetic structure with the Bruneian Malays having a distinct admixture pattern.

## Discussions

### Genomic Landscape of Bruneian Malays

The ∼5.28 million variants identified in our study have allowed, for the first time, the genomic landscape of Bruneian Malays to be explored in greater details and made comparative genomic studies against other Malay groups in the region possible. Although more than 216K of these variants have never been reported in dbSNP, the slightly lower Het/Homo ratio observed here, when compared to other population groups, seems to suggest lower heterozygosity in the genetics of the Dusun Malays than other Malay groups in the region. Higher homozygosity has in fact been observed in smaller population and less admixed groups [41], which is likely true for the Bruneian Dusun Malays. Crucially, many of the Bruneian variants were found to have minor allele frequency <0.001. This subset of rare variants is likely to be unique to the Bruneian Malays. Similar pattern has also been observed in studies assessing the allele frequency of variants in different population groups [42]. Although most of these rare variants are expected to have benign functional impacts, the existence of population-specific disease risk alleles should not be ruled out. In fact, ∼200 of the novel variants were found to be located on exonic regions of each of the Bruneian genome and majority of them are predicted to be non-synonymous which may impact genes known to be associated with genetic diseases, such as cancer, cardiovascular diseases, and diabetes mellitus, and pharmacogenetic markers. However, more genotypic and phenotypic data from larger cohorts will be needed to provide the necessary resolution power required in studies such as GWAS to establish the likely association between these rare variants and diseases.

Although majority of the variants were distributed fairly evenly across the genome, several variant-dense loci which have not been reported in other population groups were identified in the two Bruneian Malay individuals. For instance, chr10 and 13 were found to harbour the highest variant density whereas findings from 1KGP reported chr16 and 1 as having the highest and lowest SNP density, respectively [39]. However, it is important to take into consideration potential biases of such a comparison between two local individuals against the majority European samples in 1KGP.

In addition, a number of genes are predicted to be disrupted by the CNVs identified in the two local Malays. Although it may be challenging to reconstruct the exact haplotype structure of these CNVs using the data from short read sequencing, such disruptions do have the potential to produce new protein isoforms which may or may not be functional. In fact, 13 haplotypes/alleles of the *HLA-H* pseudogene were reported elsewhere to reach 19.6% in East Asian populations and they have been shown to be functional [43]. Therefore, further investigation of these CNV-impacted genes using such technology as long read sequencing may be warranted.

### Novel sequences in the Bruneian Malay genome

Although the novel sequences identified here have not been validated, for example, by Sanger sequencing or PCR, the facts that they do share high sequence identity to partial human reference genome and fosmid- or BAC-cloned human sequences have provided sufficient evidence that these are indeed human sequences. Furthermore, the shared similarity of large segments of novel sequences among the Bruneian Malay, Chinese and Japanese corroborates well with findings in our admixture analysis. Indeed, some genomic sequences which are absent in the human reference genome have now be shown to be Asian-specific and they are shared among different Asian population groups. Shi *et al*. (2016) [12] found that only a quarter of the Chinese HX1 novel sequences are absent in previously reported Asian genomes, suggesting that majority of them are likely to be present in other Asian populations. Similarly, more than half of the Japanese JRG novel sequences reported by Nagasaki *et al*. (2019) [15] were also found in the *de-novo* assembled Korean genome by Seo *et al*. (2016) [16].

Since both Chinese and Japanese studies have reported novel sequences in the Mbp range, it is probable that more Malay-specific genomic sequences have yet to be uncovered. In addition, the finding of an ORF encoding a novel human homologue of a primate’s small zinc finger protein have added more weight to the importance of finding such population-specific novel sequences.

### Subtle Genetic Differences among Southeast Asian Malays

The Malays are one of the largest Austronesian population groups spreading over Island Southeast Asia and as far away as South Africa. Although the different sub-ethnic groups share considerable cultural and linguistic ties, they are not genetically homogenous [6, 11, 44]. Adding to this genetic diversity, population genetic structure analysis conducted here has, for the first time, unveiled the differences in genetic ancestries among the Murut and Dusun Malays in Brunei and the presumably more closely related Malay groups in Singapore and Malaysia. However, both PCA and admixture analysis revealed that the local Malays actually share a closer genetic ancestry background with the Filipino Vizaya and Luzon groups than the Singaporean and Malaysian Malays. The handful Malaysians who were found to share near identical genetic ancestries with this Bruneian-Filipino group most likely belong to either the Malay or one of the indigenous groups in the Malaysian Bornean state of Sabah. In fact, Yew *et al*. (2018) [11] reported that Malays from East Malaysia, specifically those residing in the state of Sabah, share a common ancestry with the Filipinos. Given the close geographical proximity, genetic admixture among these people would not have been unexpected. Interestingly, a recent study on population genetic structure of the various Indonesian ethnic groups reported that the Sulawesians there were more closely related to the Bruneian Dusuns and Muruts than any other groups on the Indonesian Archipelago [44]. While it is likely that the Bruneian Dusun and Murut Malays may also share highly similar genetic ancestries with other yet to be studied sub-ethnic groups on the various islands of Borneo, the Philippines and Indonesia, evidence to date seems to suggest that this group of “Malay” was likely to have spread from the Philippines west- and south-ward to as far as the coastal regions of Borneo and Sulawesi.

## Conclusion

Although only two out of the seven local Malay sub-ethnic groups were included in our studies and mixed datasets consisting of WGS and genotyping were used, this is, to our best knowledge, the first and most comprehensive genetics and genomics analysis of the Malays in Brunei. In addition to adding ∼5.2 million variants to the local Malay population and the discovery of ∼227 Kbp novel genomic sequences, our studies have also shown the existence of subtle differences in the population genetic structure among the different Malay groups in Southeast Asia. Hence, a more refined stratification of these groups using variants from larger cohorts will be necessary should the benefits of medical genetics and genomics are to be fully realised in the region.

## Data availability

The consensus genetic variants for the 41 Bruneian Malays are available at: https://genome-asia.ucsc.edu/s/Mirza%20Azmi/BNMalayGRCh38.

## Acknowledgements

We are grateful to the local individual who willingly share with us his family’s genotyping data from AncestryDNA^®^.

## Author Contributions

MA designed the study, performed the bioinformatics analysis, and drafted the manuscript. AI and LC provided valuable insights and discussion on human and medical genetics. ZHL conceived and designed the study, supervised the works, and drafted the manuscript. All authors reviewed the manuscripts.

