## Supplementary Materials for "Genomic Insights of Bruneian Malays"

**Supplementary Figure Legends**

**Figure S1. Workflow for processing WGS and genotyping data of 41 Brunei Malays.** NGS reads from two individuals were first mapped against GRCh38 human reference genome before consensus variants called by three callers were obtained. Genotyping alleles were transformed to VCF file format and lifted-over to GRCh38. All variants were then merged and annotated.

**Figure S2. Workflow for discovering novel sequences in Bruneian Malays.** (A) NGS reads which did not map to the human reference genome were fed into a microbial mapping and a metagenomic analysis pipeline to remove as much microbial sequences as possible. (B) The final unmapped reads were either mapped to the Chinese/Japanese novel contigs or *de novo* assembled.

**Figure S3. Locations of mapped reads across the two Malay genomes.** More than 94% of the two genomes (blue and green bars) were covered by the NGS reads. The remaining regions were either unmapped (orange bars) or known gaps (yellow bars) in the reference genome. Large proportions of the unmapped regions fall on the centromere (red bars).

**Figure S4. BLAST alignment of an ORF derived from de-novo assembled contig against three primatal proteins.** (A) The amino acid sequence of the *de novo* assembled contig and (B) BLAST alignment between of the encoded ORF with a putative zinc finger protein of rhesus macaque.

Table S1: List of Bruneian Malay subjects analysed

| Ethnicity | Type of data | Number of subjects | | NCBI Accession No. | Original study |
|---|---|---|---|---|---|
| | | Male | Female | | |
| Dusun | Whole-genome sequencing data | - | 2 | ERX1462684, ERX1462681 | [1] |
| Dusun | Genotyping data (Illumina OmniExpress Bead Chips) | 10 | 10 | GSE77508 | [2] |
| Murut | Genotyping data (Illumina OmniExpress Bead Chips) | 7 | 10 | GSE77508 | [2] |
| Mixed-Race (Malay-European) | Genotyping data (700K SNP array) | 1 | 1 | - | AncestryDNA® |
| European | Genotyping data (700K SNP array) | 1 | - | - | AncestryDNA® |

Table S2: List of databases used in the variant annotation

| Database (URL) | Feature | Ref. |
| --- | --- | --- |
| dbSNP (https://www.ncbi.nlm.nih.gov/snp/) | A database of known short variants in the human genome | [3] |
| Ensembl gene transcripts (https://ensembl.org/) | A comprehensive database of gene transcripts for identifying the coordinate as well as the consequence of each variant | [4] |
| GnomAD (https://gnomad.broadinstitute.org/) | A global allelic frequency database for determining the relative frequency of a variant in different populations | [5] |
| ClinVar (https://ncbi.nlm.nih.gov/clinvar/) | A curated database that holds health-associated phenotypes of known human genetic variations | [6] |
| GWAS catalogue (https://ebi.ac.uk/gwas/) | A repository of GWAS | [7] |
| SIFT (http://sift-dna.org/) | A collection of predicted score for non-synonymous SNPs based on evolutionary conservation | [8] |
| PolyPhen-2 (http://genetics.bwh.harvard.edu/pph2/) | An archive of predicted score for genetic variations based on protein structure/function and evolutionary conservation | [9] |

Table S3: List of Asian populations used for the population genetic structure analysis.

| Region | Population (Country) | Population code | Sample size | Ref. |
|---|---|---|---|---|
| South Asia | Gujarati Indians (USA) | SAS_GIH | 103 | [10] |
| South Asia | Punjabi (Pakistan) | SAS_PJL | 96 | [10] |
| South Asia | Bengali (Bangladesh) | SAS_BEB | 86 | [10] |
| South Asia | Telugu Indian (UK) | SAS_ITU | 102 | [10] |
| South Asia | Tamil (UK) | SAS_STU | 102 | [10] |
| East Asia | Dai Chinese (China) | EAS_CDX | 93 | [10] |
| East Asia | Han Chinese (China) | EAS_CHB | 103 | [10] |
| East Asia | Japanese (Japan) | EAS_JPT | 104 | [10] |
| East Asia | Southern Han Chinese (China) | EAS_CHS | 105 | [10] |
| Southeast Asia | Kinh Vietnamese (Vietnam) | EAS_KHV | 99 | [10] |
| Southeast Asia | Dusun (Brunei) | BN_Dusun | 22 | [1, 2] |
| Southeast Asia | Murut (Brunei) | BN_Murut | 17 | [2] |
| Southeast Asia | Mixed Malay (Brunei) | BN_Mixed | 2 | This study |
| Southeast Asia | Malay (Malaysia) | MY_Malay | 25 | [2] |
| Southeast Asia | Malay (Singapore) | SG_Malay | 185 | [11, 12] |
| Southeast Asia | Chinese (Singapore) | SG_Chinese | 96 | [11] |
| Southeast Asia | Indian (Singapore) | SG_Indian | 83 | [11] |
| Southeast Asia | Luzon (Philippines) | PH_Luzon | 12 | [2] |
| Southeast Asia | Vizaya (Philippines) | PH_Vizaya | 4 | [2] |
| Southeast Asia | Igorot (Philippines) | PH_Igorot | 21 | [2] |
| Southeast Asia | Burmese (Myanmar) | BM_Burmese | 20 | [2] |
| Southeast Asia | Vietnamese (Vietnam) | VN_Vietnamese | 18 | [2] |
| Europe | Caucasian (Unknown) | EU_control | 1 | This study |

Supplementary Table S4. Mapping statistics of the two Malay WGS reads.

| **Summary** | **Individual 1** | **Individual 2** |
|---|---|---|
| Total number of sequencing reads | 1,531,367,144 | 1,589,109,622 |
| Reads mapped to GRCh38 plus decoy (% mapped) | 1,203,488,078 (78.6%) | 1,521,871,514 (95.8%) |
| *Mean MAPQ of mapped reads* | Q43 | Q52 |
| *Percentage of ref genome mapped* | 94.0% | 95.6% |
| *Mean coverage depth* | 36.6× | 47.0× |
| *Non-primary mapping (% of total mapped reads)** | 2,499,500 (0.2%) | 1,856,061 (0.1%) |
| *Properly mapped reads (% of total mapped reads)** | 999,539,718 (84.5%) | 1,368,612,526 (91.5%) |
| *Reads with mate mapped to a different chr (% of total mapped reads)** | 7,271,717 (0.6%) | 6,723,130 (0.4%) |
| *Singletons (% of total mapped reads)** | 4,093,031 (0.3%) | 2,670,703 (0.2%) |

* These non-primary alignments include secondary and supplementary alignments.

Table S5: Genes located within SNP-dense regions in the two Malay genomes

| Chr | Start position | SNP density | Genes within high SNP density region |
|---|---|---|---|
| **Shared by both individuals:** | | | |
| 3 | 75 Mbp | 3,248 | *FRG2C, ZNF717, ROBO2* |
| 4 | 60 Mbp | | - |
| 6 | 29 Mbp | 6,521 | *HLA* gene family |
| 8 | 3 Mbp | 6,392 | *DEF* gene family |
| 11 | 5 Mbp | 4,481 | *OR51L1, OR52J3, OR52E2, OR52A5, OR52A1, OR51V1, HBB, HBD, HBG1, HBG2, HBE1, OR51B4, OR51B5, OR51B2, OR51B6, OR51M1, OR51Q1, OR51I1, OR51I2, OR52D1, UBQLN3, UBQLNL, OR52H1, OR52B6, TRIM6, TRIM34, TRIM5, TRIM22, OR56B1, OR52N4, OR56B2P, OR52N5, OR52N1, OR52N2, OR52E6, OR52E8, OR52E4, OR52E5, OR56A3, OR56A5, OR52L1, OR56A4* |
| 11 | 26 Mbp | 3,055 | *ANO3, MUC15, SLC5A12, FIBIN* |
| 11 | 48 Mbp | 3,787 | *PTPRJ, OR4B1, OR4X2, OR4X1, OR4S1, OR4C3, OR4C5, OR4A47, TRIM51GP* |
| 16 | 5 Mbp | 2,433 | *SEC14L5, NAGPA, ALG1, EEF2KMT, RBFOX1, TMEM114, METTL22, ABAT, TMEM186, PMM2, CARHSP1, LITAFD, USP7* |
| 16 | 77 Mbp | 2,535 | *MON1B, SYCE1L, ADAMTS18, NUDT7, VAT1L, CLEC3A, WWOX* |
| 16 | 81 Mbp | 3,349 | *CMC2, CENPN, ATMIN, GCSH, BCO1, GAN, CMIP, PLCG2, SDR42E1, HSD17B2, MPHOSPH6, CDH13, HSBP1, MLYCD, OSGIN1, NECAB2, SLC38A8, MBTPS1, HSDL1, DNAAF1, TAF1C, ADAD2, KCNG4, WFDC1, ATP2C2, MEAK7, COTL1, KLHL36, USP10, CRISPLD2, ZDHHC7* |
| 17 | 21 Mbp | 2,994 | *USP22, DHRS7B, TMEM11, NATD1, MAP2K3, KCNJ12, KCNJ18* |
| 20 | 29 Mbp | 3,302 | - |
| 22 | 22 Mbp | 3,048 | *VPREB1, ZNF280B, ZNF280A, PRAME, GGTLC2, IGLL5* |
| X | 0 Mbp | 3,813 | *PLCXD1, GTPBP6, PPP2R3B, SHOX, CRLF2, CSF2RA, IL3RA, SLC25A6, ASMTL, P2RY8, AKAP17A, ASMT* |

| | | | |
|---|---|---|---|
| **Observed only Individual 1:** | | | |
| 3 | 98 Mbp | 2,455 | *GABRR3, OR5AC2, OR5H1, OR5H14, OR5H15, OR5H6, OR5H2, OR5K4, OR5K3, OR5K1, OR5K2, CLDND1, GPR15, CPOX, ST3GAL6, DCBLD2* |
| 4 | 188 Mbp | 2,404 | *ZFP42, TRIML2, TRIML1* |
| 12 | 11 Mbp | 2,526 | *PRH1, TAS2R14, TAS2R19, TAS2R31, TAS2R46, TAS2R43, TAS2R30, SMIM10L1, TAS2R42, PRB3, PRB4, PRB1, PRB2, ETV6* |
| 12 | 30 Mbp | 2,448 | *IPO8, CAPRIN2, TSPAN11* |
| 13 | 19 Mbp | 2,411 | *TUBA3C, TPTE2, MPHOSPH8, PSPC1, ZMYM5, ZMYM2* |
| 19 | 23 Mbp | 2,364 | *ZNF728, ZNF730, ZNF724, ZNF91, ZNF675, ZNF681, ZNF726* |
| **Observed only in Individual 2:** | | | |
| 10 | 56 Mbp | 2,371 | *ZWINT* |
| 14 | 106 Mbp | 2,843 | - |

Table S6: Genes located within INDEL-dense regions in the two Malay genomes

| Chr | Start position | INDEL Density | Genes within high INDEL density region |
|---|---|---|---|
| **Shared by both individuals:** | | | |
| 1 | 104 Mbp | 282 | - |
| 1 | 188 Mbp | 279 | - |
| 3 | 163 Mbp | 267 | - |
| 4 | 60 Mbp | 262 | - |
| 6 | 29 Mbp | 421 | *HLA* gene family |
| 10 | 53 Mbp | 272 | *PCDH15, MTRNR2L5, ZWINT* |
| 11 | 23 Mbp | 286 | *LUZP2, ANO3, MUC15, SLC5A12, FIBIN* |
| 11 | 50 Mbp | 305 | - |
| 11 | 98 Mbp | 263 | *CNTN5* |
| 12 | 127 Mbp | 300 | - |
| 14 | 40 Mbp | 277 | - |
| 21 | 19 Mbp | 280 | *NCAM2* |
| 21 | 37 Mbp | 282 | *RIPPLY3, PIGP, TTC3, VPS26C, DYRK1A, KCNJ6* |
| X | 0 Mbp | 413 | *PLCXD1, GTPBP6, PPP2R3B, SHOX, CRLF2, CSF2RA, IL3RA, SLC25A6, ASMTL, P2RY8, AKAP17A, ASMT* |
| **Observed only in Individual 1:** | | | |
| 4 | 28 Mbp | 261 | - |
| 7 | 64 Mbp | 266 | *ZNF722P, ZNF727, ZNF735, ZNF679, ZNF736, ZNF680, ZNF107, ZNF138, ZNF273, ZNF117, ERV3-1* |
| 18 | 71 Mbp | 273 | - |
| 19 | 23 Mbp | 294 | *ZNF728, ZNF730, ZNF724, ZNF91, ZNF675, ZNF681, ZNF726* |
| **Observed only in Individual 2:** | | | |
| 8 | 30 Mbp | 262 | *SARAF, LEPROTL1, MBOAT4, DCTN6, RBPMS, GTF2E2, SMIM18, GSR, UBXN8, PPP2CB, TEX15, PURG* |
| 10 | 60 Mbp | 259 | *ANK3, CDK1, RHOBTB1* |

Table S7: List of protein-coding genes within CNVs.

| **Chr** | **Start** | **End** | **CNV size (Kbp)** | **Gene affected** | **Region of gene** | **Associated disease** | **CN1*** | **CN 2*** |
|---|---|---|---|---|---|---|---|---|
| 1 | 103,623,000 | 103,626,000 | 3 | *AMY2A* | Intron7-3'UTR | Obesity [13] | 3 | 2 |
| 1 | 109,686,000 | 109,702,000 | 16 | *GSTM1* | 5'UTR-3'UTR | Cancer [14] | 0 | 2 |
| 1 | 145,375,000 | 145,379,000 | 4 | *NBPF20* | intron28-intron33 | Cancer [15] | 2 | 1 |
| 1 | 196,822,000 | 196,837,000 | 15 | *CFHR1* | intron1-3'UTR | Macular degeneration [16] | 2 | 3 |
| 4 | 68,510,000 | 68,625,000 | 115 | *UGT2B17* | 5'UTR-3'UTR | Osteoporosis [17] | 0 | 2 |
| 6 | 29,884,000 | 29,938,000 | 54 | *HLA-H*[#] | 5'UTR-3'UTR | Immune responses [18] | 2 | 0 |
| 6 | 160,633,000 | 160,636,000 | 3 | *LPA* | intron6-intron8 | Coronary artery disease [19] | 6 | 0 |
| 9 | 678,590,00 | 678,650,00 | 6 | *ANKRD20A1* | exon1-intron3 | Cancer [20] | 0 | 2 |

[*]Number of CNV in individual 1 and 2.

[#]Variants of HLA-H pseudogenes have been to encode full-length gene.

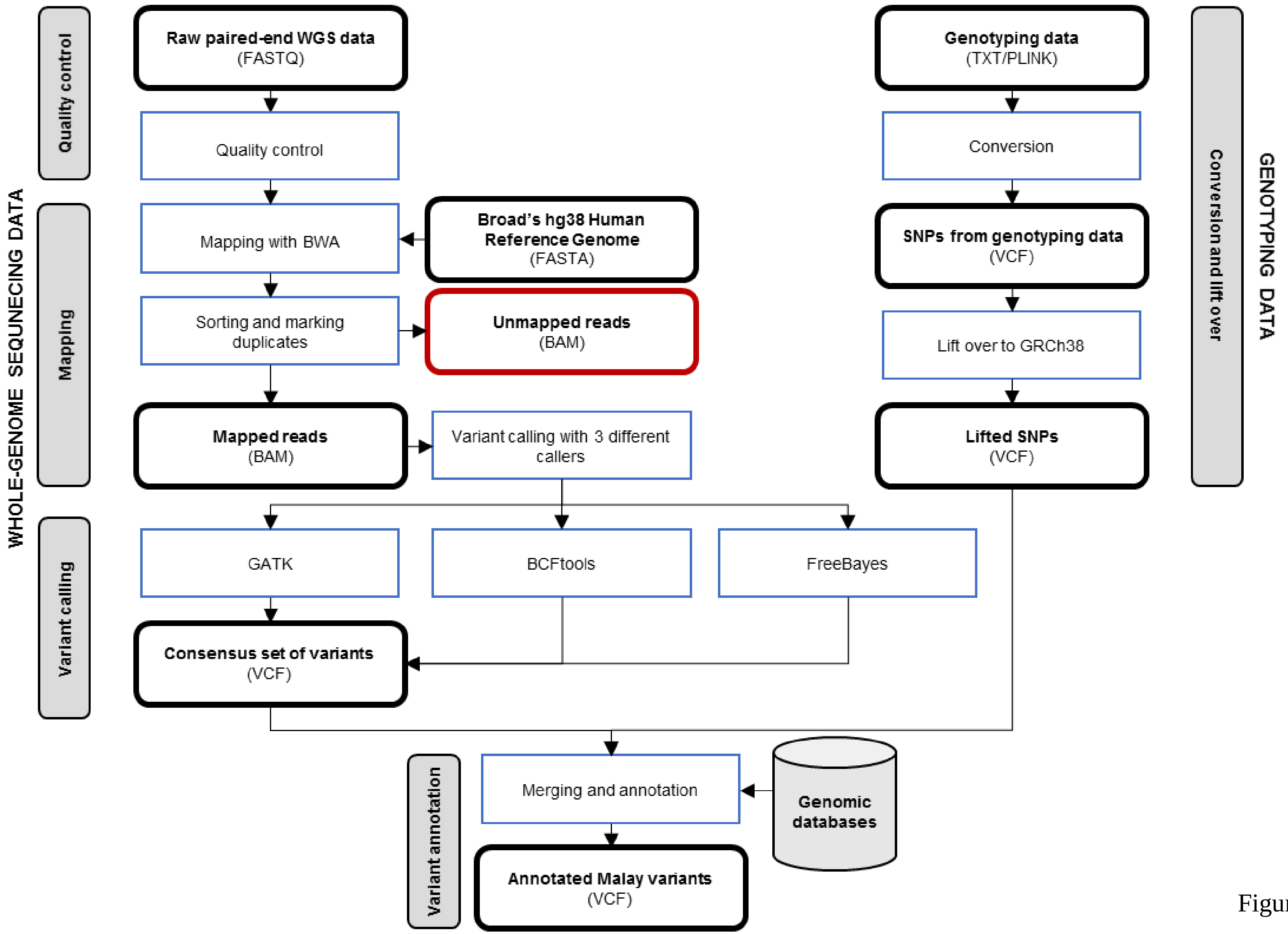
WHOLE-GENOME SEQUNECING DATA
Quality control
Mapping
Variant calling
Raw paired-end WGS data
(FASTQ)
Quality control
Mapping with BWA
Broad's hg38 Human Reference Genome
(FASTA)
Sorting and marking duplicates
Unmapped reads
(BAM)
Mapped reads
(BAM)
Variant calling with 3 different callers
GATK
BCFtools
FreeBayes
Consensus set of variants
(VCF)
Genotyping data
(TXT/PLINK)
Conversion
SNPs from genotyping data
(VCF)
Lift over to GRCh38
Lifted SNPs
(VCF)
Conversion and lift over
GENOTYPING DATA
Variant annotation
Merging and annotation
Genomic databases
Annotated Malay variants
(VCF)

Figure S1

(A)

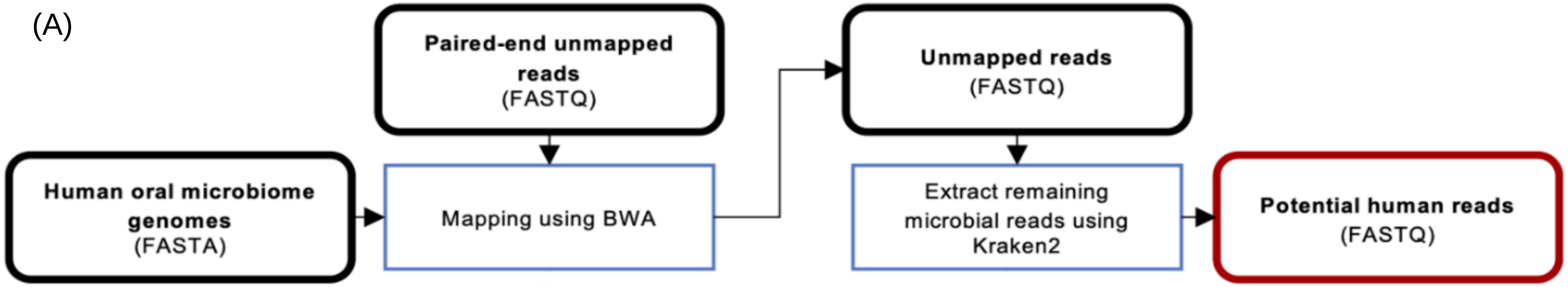
Paired-end unmapped reads (FASTQ)
Unmapped reads (FASTQ)
Human oral microbiome genomes (FASTA)
Mapping using BWA
Extract remaining microbial reads using Kraken2
Potential human reads (FASTQ)

Figure S2A

(B)

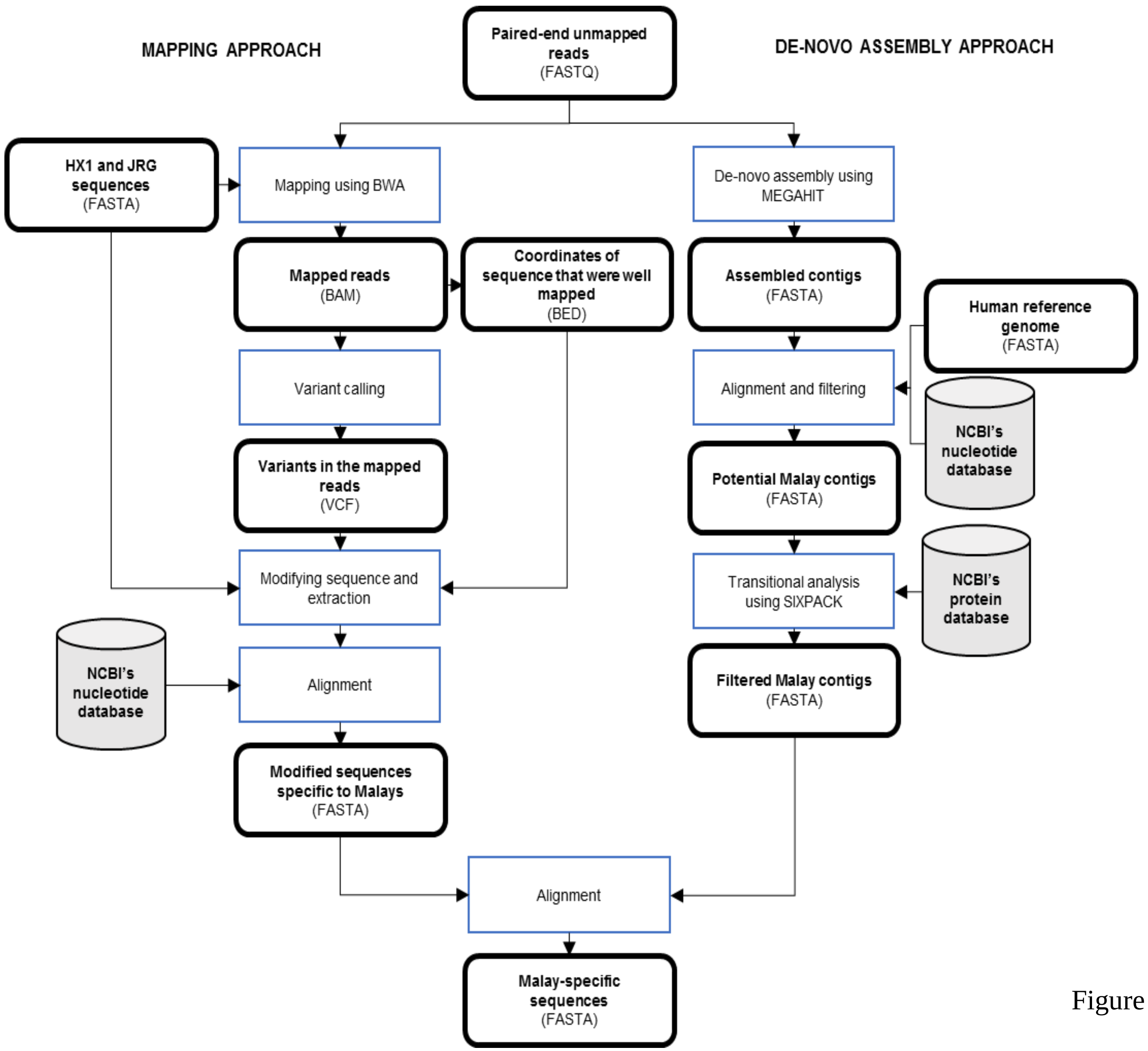

MAPPING APPROACH
Paired-end unmapped reads (FASTQ)
DE-NOVO ASSEMBLY APPROACH
HX1 and JRG sequences (FASTA)
Mapping using BWA
De-novo assembly using MEGAHIT
Mapped reads (BAM)
Coordinates of sequence that were well mapped (BED)
Assembled contigs (FASTA)
Human reference genome (FASTA)
Variant calling
Alignment and filtering
NCBI's nucleotide database
Variants in the mapped reads (VCF)
Potential Malay contigs (FASTA)
Modifying sequence and extraction
Transitional analysis using SIXPACK
NCBI's protein database
NCBI's nucleotide database
Alignment
Filtered Malay contigs (FASTA)
Modified sequences specific to Malays (FASTA)
Alignment
Malay-specific sequences (FASTA)


Figure S2B

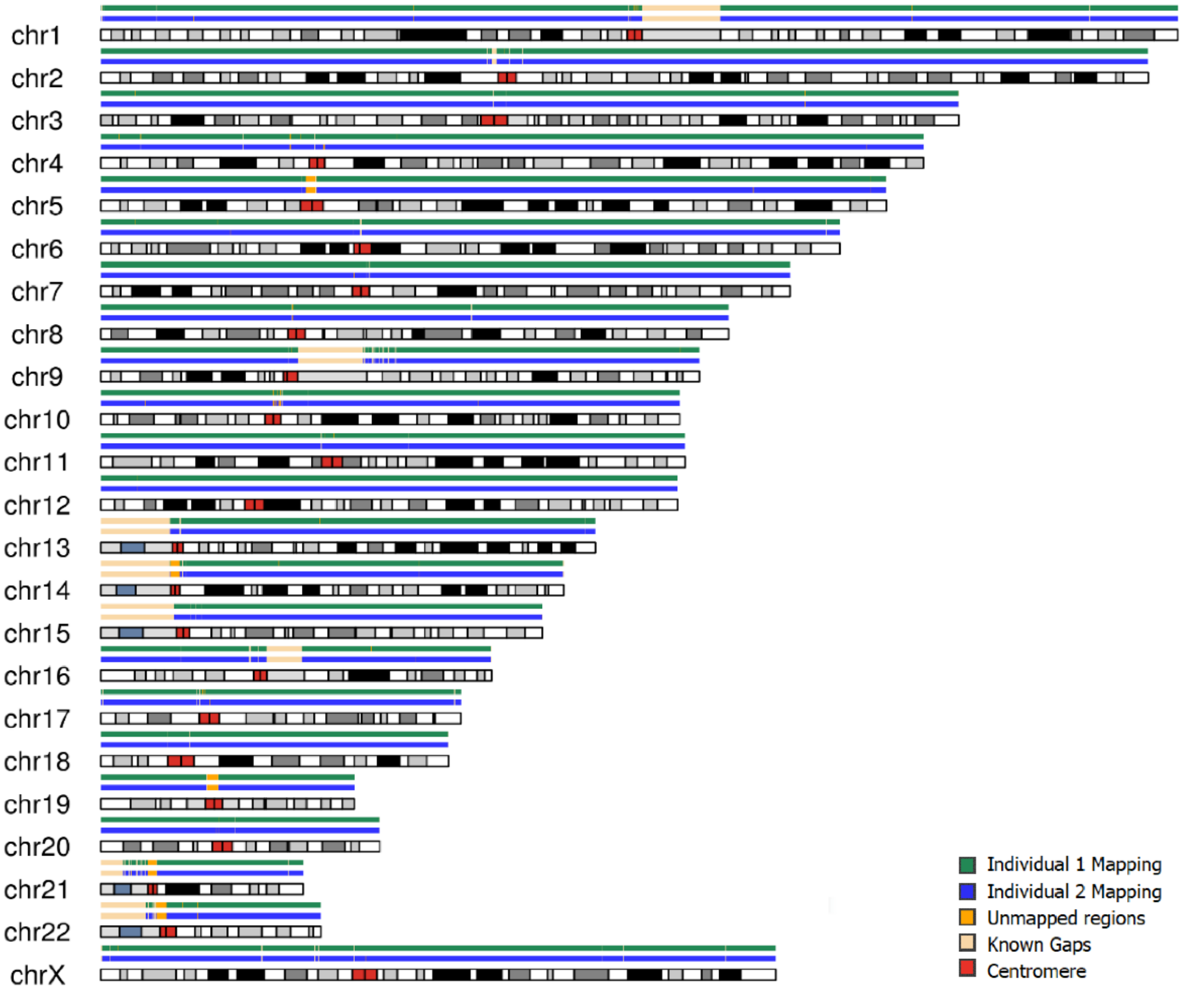
chr1
chr2
chr3
chr4
chr5
chr6
chr7
chr8
chr9
chr10
chr11
chr12
chr13
chr14
chr15
chr16
chr17
chr18
chr19
chr20
chr21
chr22
chrX
Individual 1 Mapping
Individual 2 Mapping
Unmapped regions
Known Gaps
Centromere

Figure S3

```
A    >Translation of k31_201675 in frame 3, ORF 3, threshold 50, 159aa
     LKHQHVQTGGKPYVYPQHGKTFSVNPSPMKQQRSHPGEDLHESKECGKIFRHWILSDIRELLWISSVWKAFCAS  75
     SVLLNISKFILDKSLIYIREYSKASKCCDSLIKHQRICTGEKPCWSEEDSKRFIMVQLWSPTRESTLESSTIMN  150
     VVCEGRPLFGS                                                                 159


B    >EHH18077.1 hypothetical protein EGK_14614 [Macaca mulatta]
     Identities = 97/106 (92%), Positives = 100/106 (94%), Gaps = 0/106 (0%)

     ORF         54   ILSDIRELLWISSVWKAFCASSVLLNISKFILDKSLIYIREYSKASKCCDSLIKHQRICT  113
                      +LSDIRELLWISSVWKAFC SSVLLNISKFILDKSLIYIR  SKASKCCDSLIKHQRICT
     M.MULATTA   13   VLSDIRELLWISSVWKAFCVSSVLLNISKFILDKSLIYIRGCSKASKCCDSLIKHQRICT  72


     ORF         114  GEKPCWSEEDSKRFIMVQLWSPTRESTLESSTIMNVVCEGRPLFGS  159
                       EKP WSEEDSKRFI+VQLWSPTRESTLESST+MNVV EGRPLFGS
     M.MULATTA   73   AEKPYWSEEDSKRFIVVQLWSPTRESTLESSTVMNVVYEGRPLFGS  118
```

Figure S4
